# Technical Note: A fast and robust integrator of delay differential equations in DCM for electrophysiological data

**DOI:** 10.1101/2020.12.28.424540

**Authors:** Dario Schöbi, Cao-Tri Do, Stefan Frässle, Marc Tittgemeyer, Jakob Heinzle, Klaas Enno Stephan

## Abstract

Dynamic causal models (DCMs) of electrophysiological data allow, in principle, for inference on hidden, bulk synaptic function in neural circuits. The directed influences between the neuronal elements of modeled circuits are subject to delays due to the finite transmission speed of axonal connections. Ordinary differential equations are therefore not adequate to capture the ensuing circuit dynamics, and delay differential equations (DDEs) are required instead. Previous work has illustrated that the integration of DDEs in DCMs benefits from sophisticated integration schemes in order to ensure rigorous parameter estimation and correct model identification. However, integration schemes that have been proposed for DCMs either emphasise speed (at the possible expense of accuracy) or robustness (but with computational costs that are problematic in practice).

In this technical note, we propose an alternative integration scheme that overcomes these shortcomings and offers high computational efficiency while correctly preserving the nature of delayed effects. This integration scheme is available as open-source code in the Translational Algorithms for Psychiatry-Advancing Science (TAPAS) toolbox and can be easily integrated into existing software (SPM) for the analysis of DCMs for electrophysiological data. While this paper focuses on its application to the convolution-based formalism of DCMs, the new integration scheme can be equally applied to more advanced formulations of DCMs (e.g. conductance based models). Our method provides a new option for electrophysiological DCMs that offers the speed required for scientific projects, but also the accuracy required for rigorous translational applications, e.g. in computational psychiatry.

## Introduction

A key goal of Translational Neuromodeling (TN) and Computational Psychiatry (CP) is the development of generative models as “computational assays” and their application to clinical questions in psychiatry (Stephan and Mathys, 2014). One branch of the development of computational assays concerns circuit models which represent distinct aspects of synaptic function, such as different types of receptors and neuromodulatory processes, at the level of interacting neuronal populations. By inverting these models using Bayesian techniques, the goal of TN/CP is to identify subject-specific alterations of synaptic function which may support the stratification of heterogeneous disorders and individual treatment predictions (Stephan et al., 2015). For example, computational assays of this sort have been used to examine dysfunction of specific ion channels in monogenetic channelopathies (Gilbert et al., 2016) or alterations of NMDA receptor function due to autoimmunological processes (Symmonds et al., 2018).

One candidate method for this purpose is Dynamic Causal Modelling (DCM) for electrophysiological measures (David et al., 2006; Moran et al., 2008; Moran et al., 2013) as implemented in the open-source software SPM (https://www.fil.ion.ucl.ac.uk/spm/). Due to the high temporal resolution of electroencephalography (EEG) or magnetoencephalography (MEG), data acquired with these methods contain information about (average) synaptic processes in (large) neuronal populations (Gilbert et al., 2016; Neymotin et al., 2020). DCM for EEG/MEG models electrical scalp potentials or magnetic fields, respectively, as arising from the dynamics of large neuronal populations (neural masses) that interact which each other through short-range and long-range synaptic interaction. The latent dynamics are based on the flow of charged ions across the cell membrane during the generation of action potentials, dependent on the density of afferent axonal connections, the density of receptors in the membrane and (indirectly) neurotransmitter availability. These synaptic actions then describe the dynamics of a population of cell types within a cortical column. In the simplest variant of DCM for EEG, two distinct populations of excitatory cell types (stellate and pyramidal cells) and one inhibitory population constitute a single cortical column. These populations are intrinsically (within-column) connected to each other, and extrinsically connected to different columns. Importantly, due to the finite transmission speed of axonal connections, the directed influence one population exerts over another is subject to delays. These delays render the differential equations used to describe the neuronal population dynamics delay differential equation (DDEs), requiring specialised integration schemes.

The current default integration scheme employed for electrophysiological DCMs in SPM is based on an extension of an integration scheme for ordinary differential equations (ODEs) ((Ozaki, 1992), for review, see (Ostwald and Starke, 2016)). This scheme effectively absorbs delays into an adaptive, delay-dependent step size of a standard ODE step and is computationally highly efficient. This computational efficiency is important since parameter estimation through approximate Bayesian inference techniques (e.g. variational Bayes or Markov chain Monte Carlo) typically requires a large number of integrations.

In this technical note, we show simulations, some based on examples with analytical solutions, where this particular DDE scheme fails to account for delayed effects accurately, causing non-negligible errors. This extends previous work by Lemarechal and colleagues (Lemarechal et al., 2018) who demonstrated that, in DCM for EEG, model selection and parameter inference can be affected considerably by the choice of integration method and that the DDE integration scheme in SPM may not always be highly accurate. Lemarechal et al. (2018) also demonstrated that alternative DDE integration schemes (i.e. the *dde23* integrator in MATLAB) provided accurate estimates of DCM parameters, however, at the cost of impractically long compute times.

Here, we extend the work by Lemarechal and colleagues. by presenting an integration scheme newly applied to these models – Linearized Delayed Euler (*LDE*) – that combines robustness with computational efficiency^1^. Using simulated and empirical data, we show that this method accurately accounts for delayed effects but is orders of magnitude faster than the robust DDE integration method proposed by Lemarechal and colleagues. Our DDE integration scheme is compatible with SPM and freely available as part of the open-source software package TAPAS (https://www.translationalneuromodeling.org/tapas).

## Methods

In general, 1^st^ order delay differential equations (DDEs) are characterized by the following set of equations

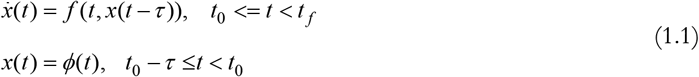

where *x* denotes the state variable or state, *t* denotes time and *t*_0_ and *t*_*f*_ the initial and final timepoint, respectively. The initial state function is defined by *ϕ*, and *τ* ∈ [0, ∞) are semi-positive delays (Bellen and Zennaro, 2013).

Introducing delays can have a number of important implications for the behavior of a system. Most prominently, delays can stabilize or destabilize a system, can lead to non-uniqueness of solutions, and may make a system exhibit oscillatory or chaotic behavior when compared to the sibling ODE system (Bellen and Zennaro, 2013). One method to deal with simple delay problems uses a fine-grained integration mesh such that all delayed states fall on grid-points. An integration step can then be computed via a classical ODE step, e.g. the forward Euler method (Elsgolts, 1964). In his doctoral thesis, Feldstein (1964) introduced a method where the grid-points are independent of the delays. He combined an Euler step with a linear interpolation between grid-points to approximate delayed states (Equation 3.1.3 in (Bellen and Zennaro, 2013)):

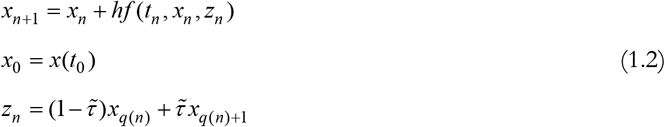

Here *q*(*n*) and *q*(*n*) + 1 code for the timepoints immediately preceding and succeeding the delayed time, 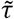 the delayed time normalized to this interval and *h* the integration step size. Hence, *z*_*n*_ is simply the piecewise linear approximation on *x*(*t* − *τ*). Because of the freedom in the choice of interpolation and the discrete ODE step, the previous equations are merely an example of a whole class of DDE integration schemes, the so-called *continuous extension* of ODE methods.

In this technical note, we will evaluate a continuous extension of ODE method, in combination with a forward Euler step, for the particular problem of integrating DDEs in DCM. For a more comprehensive list of DDE integration methods, we refer to existing literature (e.g. (Balachandran et al., 2009; Erneux, 2009; Smith, 2011; Bellen and Zennaro, 2013)).

### Local linearization delayed integration (*spm_int_L*)

In SPM12 (version 6906), the commonly used integration scheme (*spm_int_L*) for the integration of a system of dynamical equations is based on developments by Ozaki (1992). Under the assumption that the system is locally linear (in time), the integration can be performed efficiently and robustly through an adaptive step size incorporated in the Jacobian 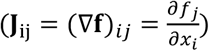 (the derivation of this update is provided in Ozaki’s 1992 article):

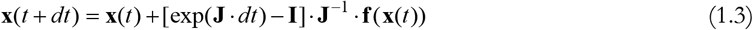

Here, **I** denotes the identity matrix. The proposed solution for DDEs, *spm_int_L*, implements delays efficiently but approximately in the Jacobian, which is motivated from a second linearization of the following form (Ostwald and Starke, 2016):

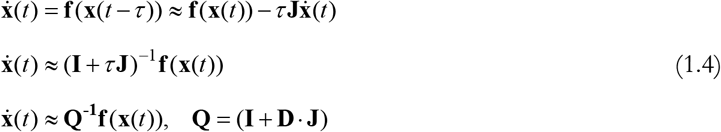

In higher-dimensional problems, the constant delay *τ* turns into a delay matrix **D**. In the following, we will refer to this integrator as *spm_int_L* (i.e. the name of the function implementing this scheme in SPM12). Combining the last line of Eq. (1.4) with Eq. (1.3), the update equations for *spm_int_L* are as follows:

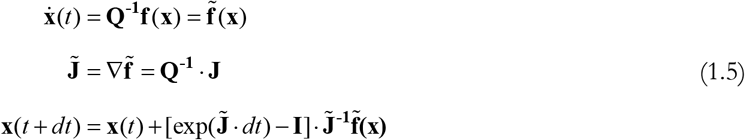

and therefore

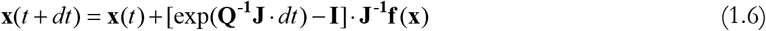

The advantage of this integration scheme is that one does not need to keep track of the history of the states, as **f**(**x**(t)) is only ever evaluated at the current timepoint. This makes the approach very efficient. Also, if the system is linear in the states (an additional assumption by *spm_int_L*, despite the non-linearity due to a sigmoid transformation), one only needs to compute the Jacobian once since ∇**f** is constant. There are schemes in place which would allow for the continuous evaluation of the Jacobian (see *spm_int_J*.*m*). However, they render integration one order of magnitude slower. A crucial question about this method is how well the 1^st^ order Taylor approximation on the dynamical equations (Eq. (1.4)) performs, particularly for longer delays of extrinsic connections.

### Continuous extensions of ODE methods and Linearized Delayed Euler (*LDE*)

Euler (forward) integration schemes have been around for two centuries and assume that for a small timestep *dt*, the system will evolve in the direction of its gradient (in time), i.e.

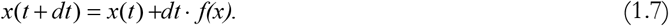

Obviously, delays are not incorporated in Eq. (1.7). In the spirit of continuous extension of ODEs (Feldstein, 1964), we used an approximation to the states 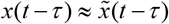 by linearly interpolating between the two neighboring, evaluated timesteps:,

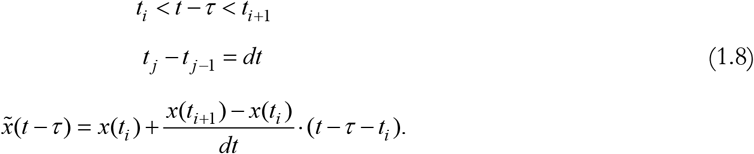

In full, the update (*φ*) of the Linearized Delayed Euler scheme (*LDE*) is given by

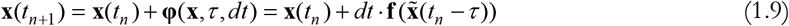

A graphical illustration is provided in **Figure 1**. Please note that we used a shorthand notation here. States **x** denote state vectors and, analogously, the delays *τ* = *τ*_*ij*_ are state-dependent. Therefore, Eq. (1.9) is an extension of Eq. (1.8) for a system of DDEs with state-dependent delays.

**Figure 1.**
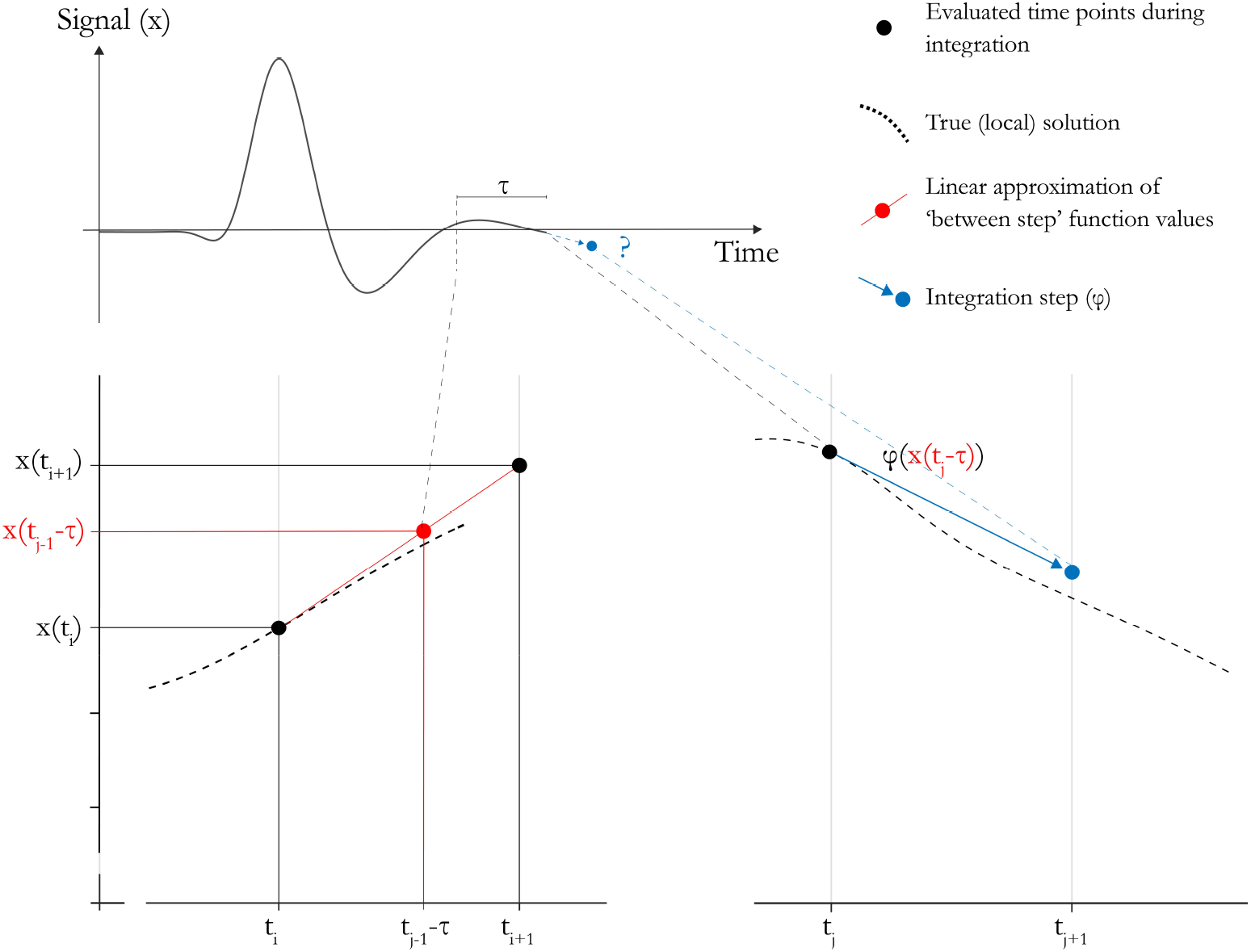
Schematic overview of the implementation of delays. The update function ***φ*** requires function values at previous timepoints. The function at times between sampled timepoints is linearly approximated for efficiency. The update step is an arbitrary ODE step (e.g. Euler).

Two major points need to be considered for the choice of interpolation method: one is the *propagation of discontinuities* (resulting from jumps in the derivative at the transition point), the other one concerns so-called *overlapping*, i.e. when delays become smaller than the step size of integration. The second problem can be easily solved by reducing the step size. The first problem is more difficult; however, it is not critically relevant in the context of DCM as the solution can be expected to be relatively smooth at transition points. This is because in the typical application scenario of modeling cortical circuits in mammalian brains, the lag of the driving input is longer than the delays within the circuit. In this case, the initial conditions are a steady state of the system.

One intriguing property here is that the continuous extension 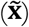 is independent of the discrete ODE step (**φ**) used, and the same linear approximation of delayed states could be combined with a different method of updating. For simplicity, we omit comparisons beyond simple forward Euler updates. A broader overview on these topics and extensions, using for example the Ozaki update in combination with the continuous extension, are provided in (Schöbi, 2020).

### Simulations

We simulated responses based on three simple (delayed) dynamical systems. The systems were chosen for different reasons: for the first example, there exists the particular case of a delay magnitude with an analytical solution that allows for clear predictions about the qualitative properties the integrated signal should display. The 2^nd^ and 3^rd^ example approximate and equal, respectively, the complexity of the dynamical system underlying the convolution-based DCMs for event-related potentials (ERPs); this model is explained below. For all systems, we parametrically changed the magnitude of the delay to assess the regime of numerical stability of the approximations used in the respective integration scheme. To this end, we compared the two integration schemes mentioned above – *spm_int_L* and *LDE* – to an integration scheme for DDEs in MATLAB, *dde23*, which is Runge-Kutta based and allows for the definition of local error bounds (Shampine et al., 2000; Shampine and Thompson, 2001). Lemarachal and colleagues (2018) also used *dde23* as a reference. All simulations were performed at an integration step size of 1 ms. For *dde23*, we used a local error tolerance of 1E-6.

#### Delayed, one-dimensional exponential decay

Consider the following dynamical system (**Figure 2A**)

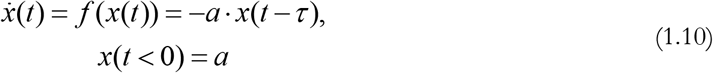

**Figure 2.**
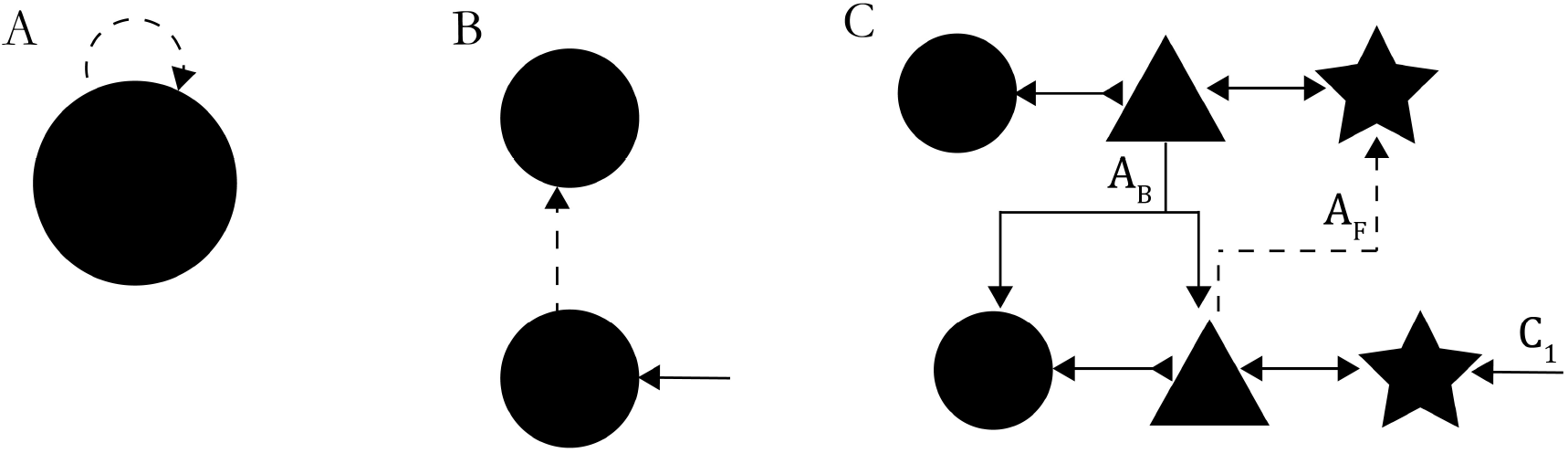
Simulation setup of delay differential equations for the three dynamical systems. Dotted lines represent delayed connections. A) One-dimensional exponential decay; B) Two damped, harmonic oscillators with forward coupling; C) Convolution-based DCM for ERP. Symbols depict distinct neuronal populations; arrowheads depict inhibitory / excitatory influence. Layer specificity is not shown.

While simple, this system has an interesting property. For *τ* = 0, it is simply a decaying exponential *x*(*t*) = *a* · *exp(*−*at*). For *τ*= *π*/2, the analytical solution (except for short-lived effects introduced by the initial condition) is a superposition of a sine and cosine function, i.e. an oscillating system. Thus, we would assume that as the delays are increased, the system will start to show oscillatory behavior.

Under the approximations of *spm_int_L* the integration steps are given by

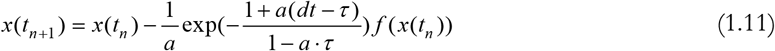

In the one-dimensional case, *Q* = (1 + *D · j*) (as per Eq. (1.4)) is a scalar, and therefore, the *spm_int_L* update is equivalent to a forward Euler update of the non-delayed system, with an adjusted step size of 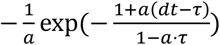. Hence, if the system is integrated with *spm_int_L*, it will never show oscillatory behavior. In addition, for finite *dt*, the system shows a singularity for *τ*= *1*/*a*.

#### Two damped, harmonic oscillators with forward coupling

For the second example, we considered two coupled harmonic oscillators (HOs), a system that closely resembles the equation underlying convolution-based DCMs:

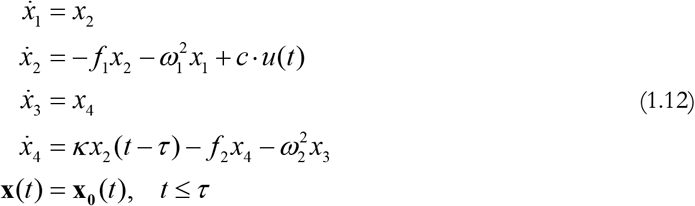

Here, the second HO receives delayed (by *τ*) input with coupling strength *κ* from the first (**Figure 2B**). The other variables parameterize the HO. Initial conditions are defined by **x**_0_(*t*). Only the first oscillator is driven by some external driving input *u*(t). All obvious dependencies on t are omitted. This system is in close analogy to the dynamics of two coupled neuronal populations of the DCM for ERP formalism. The only difference arises from constraints on the parameter (e.g. critical damping, equality of parameters across oscillators, etc.) and the lack of a sigmoid transformation in the output of the populations (effectively a sigmoid transformation of state *x*_2_ in the coupling to *x*_4_). In other words, this system constitutes a minimal set of a DCM, with only two populations and one connection.

#### Convolution-based DCM for ERP

Finally, we turn to the case of a basic convolution-based DCM with three populations. This model, originally introduced by David et al (2006), can be applied to different features of electrophysiological recordings. Here, we consider its application to ERPs (i.e. averaged responses over trials).

We connected two regions, with a forward connection from region 1 to region 2, and a backward connection from region 2 to region 1 (**Figure 2C**). Region 1 received a standard, truncated Gaussian driving input (the default input in the DCM for ERP framework (see *spm_erp_u*.*m*)). This input excites the network, resulting in a change of the post-synaptic potentials *v*(*t*) of the populations over time, modeled as

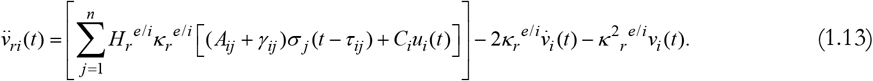

These equations come from a convolution operation between incoming, pre-synaptic firing (*σ*) of other populations and a convolution kernel (*h*)

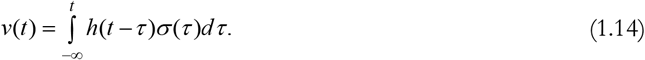

This convolution kernel *h*(*H, κ, A, γ, C*) incorporates the joint effects of the synaptic kernel *H*, the connection parameters *A, γ* and the input parameters *C*; it reflects the overall synaptic responses of a population to incoming inputs. In Eq. (1.13), the subscripts *i* and *r* refer to the *i*^th^ population of region *r*, the superscripts (*e*/*i*) indicate the excitatory or inhibitory nature of a connection. The driving input *u*(t) defines the exogenous stimulation.

Importantly, the pre-synaptic firing of a population results in a delayed (indicated by *τ*_*ij*_) de-/hyper-polarization of a connected population, hence posing the problem of integrating delayed differential equations. The transformation from post-synaptic potential to synaptic firing undergoes a sigmoid transformation (*σ*), rendering the system in Eq. (1.13) non-linear.

## Results

### Integration of 1D, HO and ERP system

To illustrate the general behavior of the two integration schemes outlined above and to assess the representation of delays, we started with two simple dynamical systems where the effects of delays can be understood analytically (first example) or intuited (second example). As a third example, we integrated a convolution-based DCM for ERPs consisting of two reciprocally connected sources. The specification of the parameters, delays, and initial conditions for the integration of the three systems are provided in **Table 1** (the corresponding dynamical equations are specified in Eq. (1.10), (1.12) and (1.13)). For the decaying exponential, we chose a decay constant such that the signal decays reasonably fast over the integration interval. For the HO, we opted for a parametrization such that the outputs of the oscillators show qualitatively similar oscillations as under standard DCM for ERP assumptions. Finally, for the full DCM model, we chose a strong forward connection (*A*_*F*_), a backward connection (*A*_*B*_) set to average strength, a driving input entering only the first region (*C*_1_) and a delay from first to second region (**Figure 2C**). Put simply, this corresponded to a higher-dimensional sibling of the harmonic oscillator simulations (second example).

**Table 1.** Integration settings for the delayed, one-dimensional exponential decay (1D), the coupled harmonic oscillators (HO) and the convolution-based DCM (ERP). All parameters not explicitly stated for the ERP model were set to the values of the prior means in SPM’s procedure for parameter estimation. The values for the parameters of the ERP model correspond to their natural description in log-space. Time window of integration ***t*** ∈ [**0, 0. 5**] ***s*** and step size ***dt*** = **0. 001 *s*** were kept equal for all systems.

| System | Initial condition | Parameters | Delays |
| --- | --- | --- | --- |
| 1D | $x(t < 0) = 10$ | $a = 10$ | $\tau \in [0, 0.1]$ |
| HO | $\mathbf{x}(t < 0) = \begin{bmatrix} 0 \\ \vdots \\ 0 \end{bmatrix}$ | $w_1 = 10 \cdot \pi$<br>$w_2 = 10 \cdot \pi$<br>$f_1 = 20$<br>$f_2 = 20$<br>$\kappa = 6 \cdot \pi$ | $\tau \in [0, 0.1]$ |
| ERP | $\mathbf{x}(t < 0) = \begin{bmatrix} 0 \\ \vdots \\ 0 \end{bmatrix}$ | $A_F = 1$<br>$A_B = 0$<br>$C_1 = 0$ | $D_{1 \rightarrow 2} \in [0, 1.6]$ |

When inspecting the integrated signals in **Figure 3**, one can observe that the linear approximation in *spm_int_L* results in three, non-negligible types of errors with regard to the qualitative and quantitative effects that delays impose on the dynamics.

**Figure 3.**
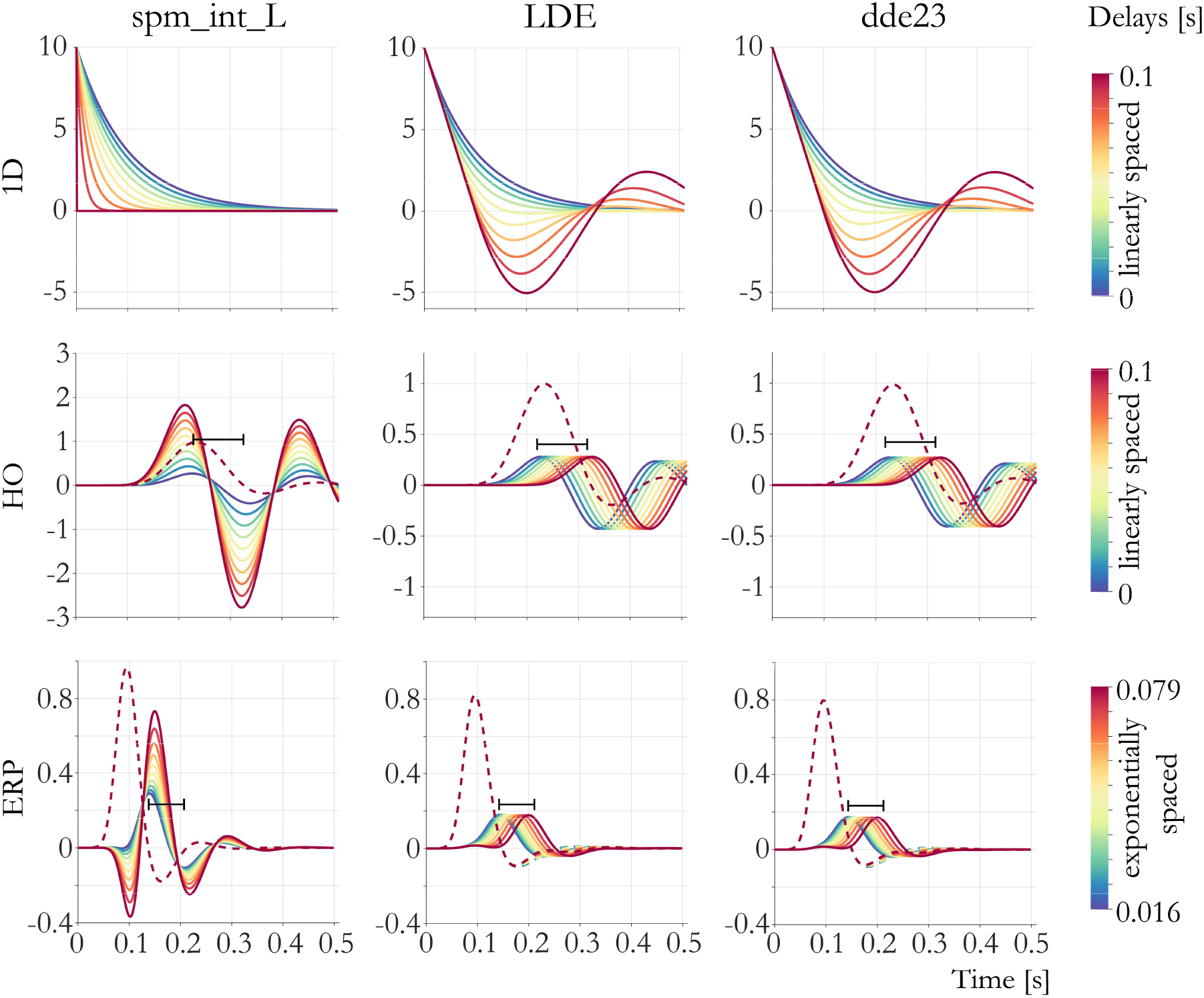
Integrated signals for different levels of delays for all three systems (top: exponential decay, middle: coupled HO, bottom: ERP). MATLAB’s RK-based integration scheme (*dde23*) taken as a reference. The black bar illustrates the maximum delay. For the coupled HO and the ERP model, the dotted line illustrates the response of the first region (not directly subject to delays). For visual clarity, for the ERP model we only show voltage traces of pyramidal cells. The dotted lines represent the activity of the non-delayed (i.e. driven) oscillator / populations.

First, and most strikingly, in the case of the one-dimensional delayed exponential decay, it fails to produce the oscillations that are predicted by the analytical solution for this system (see Methods). Instead, it reaches a mathematical singularity (i.e. infinitely fast exponential decay towards zero) at delays equal to the time constant of the system.

Additionally, *spm_int_L* violates predictions about the temporal succession of activity within the network for the dynamics of a coupled HO and the ERP model. Activity in the second (delayed) region in both systems is evoked at times forbidden under the increasing delays (its activity starts deviating from zero at times shorter than the delay), violating causality (see the length of the black bar in comparison with activity evoked under the two extreme delay conditions). Also, an artefactual peak arises in the ERP model at around 100 ms. This also emerged in the simulations by Lemarechal et al. (2018). In addition, changes in delays lead to large effects on the amplitude in the second regions, but hardly impact peak times.

Finally, the approximation fundamentally affects the underlying frequencies represented in the system (as is visible in the period of the oscillations in **Figure 3** or in a Fourier transform of the signal (not shown)).

On the other hand, the continuous extension for ODE methods proposed in the present paper, i.e. the *LDE* scheme, performs well in all three settings. The error remains small even for the delays of highest magnitude, preserving the frequencies of the true systems. Provided a reasonably small step size is used and delays do not exceed critical thresholds ^2^, the new scheme thus represents a viable alternative to the computationally more expensive reference scheme *dde23*.

We formally assessed the delay-dependent integration error by computing the difference as well as the Pearson correlation coefficient of the integrated signals compared to the reference scheme (*dde23*). The results are summarized in **Figure 4**. One can relate the integration errors to the intrinsic time constants of the system. Evidently, *spm_int_L* performs reasonably well if delays are small wrt. the time constants of the system: as can be seen in the rightmost column of **Figure 4**, the correlation between the integrated signal by *spm_int_L* and the reference *dde23* is very high for small delays and only drops to *ρ* = 0.8 for delays of around *τ* = 80, 15 and 27 ms for the three systems; these delay magnitudes correspond to approximately 80%, 6% and 13.5% of the time constants (**Table 2**). By contrast, *LDE* showed stable performance (near-perfect correlation with *dde23*) across all delays. We will discuss these implications for real data in the Discussion.

**Table 2.** Underlying frequencies, intrinsic time constants and critical delays where the Pearson correlation between *spm_int_L* and *dde23* drops to 0.8. The dominant frequencies were computed through a Fourier transform. Please note that in the case of the HO and ERP system, the frequency components of the signal lie in some range around the dominant frequency.

| System | dominant intrinsic frequencies | intrinsic time constant (in ms) | $\tau(\rho = 0.8)$ |
| --- | --- | --- | --- |
| 1D | N.A. | 100 | 80 (80 %) |
| HO | $\sim 4$ Hz | $\sim 250$ | 15 (6 %) |
| ERP | $\sim 5$ Hz | $\sim 200$ | 27 (13.5 %) |

**Figure 4.**
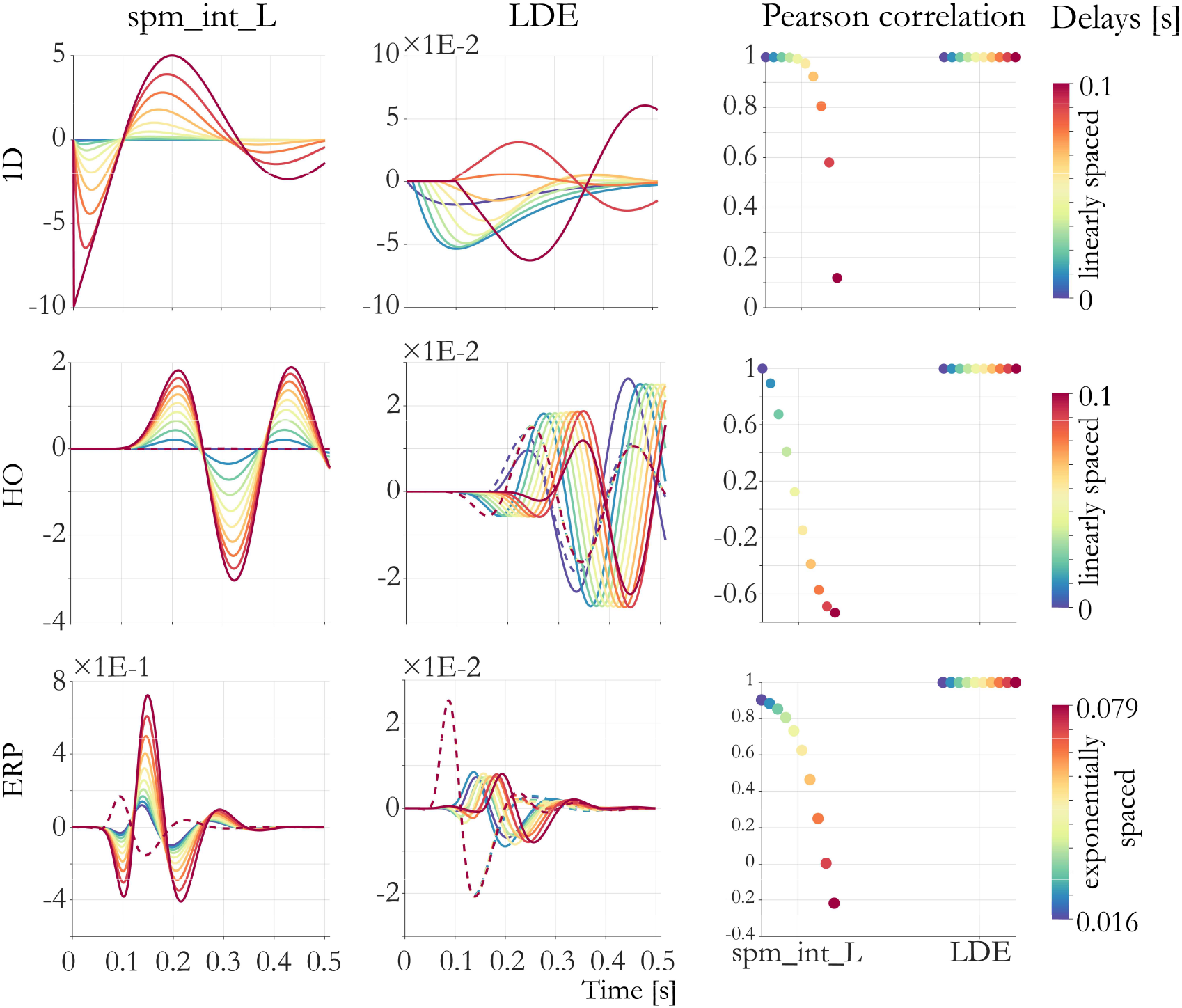
Integration error of *spm_int_L* (left column) and *LDE* (middle column) scheme for different levels of delays for all three dynamical systems. MATLABs RK-based integration scheme (*dde23*) taken as a reference. Right column: Pearson correlation coefficient of the integrated signals obtained by *spm_int_L* and *LDE*, respectively, with signals obtained by *dde23* for all delays. For visual clarity, the ERP model only shows voltage of pyramidal cells. Please note the difference in the scaling of the errors.

### Application to empirical data

The profound differences observed in our simulations suggest that the choice of integration scheme for DDEs matters when using DCM for its typical purpose, i.e., estimating the parameters of a circuit from empirical data using Bayesian inference and identifying the most plausible circuit structure through Bayesian model selection. This issue has already been investigated by (Lemarechal et al., 2018); here, we extend this analysis by comparing the parameter estimates and model selection results from an existing dataset (Jung, 2013; Jung et al., 2013) between the classical *spm_int_L* and the newly proposed *LDE* integration schemes.

In brief, we inverted a two-region-DCM^3^ of local field potential (LFP) measures in rats from primary (A1) and secondary auditory cortices (PAF). All animals (*N*=8) underwent an auditory mismatch negativity paradigm (MMN), and we investigated how the occurrence of an unexpected (deviant) tone altered the connectivity between these two auditory sources (for a complete DCM analysis based on the new *LDE* integration scheme, see (Schöbi et al., 2020)). Thus, the network used was very similar to the one illustrated in **Figure 2C**, with the exception that we used a four cell-population model (canonical microcircuit) to model a single cortical column (Bastos et al., 2012). We included all different options how a deviant might change either synaptic gain of the regions, or the strength of long range, extrinsic (presumably glutamatergic) connections, resulting in a model space of 2^4^ = 16 models. For each model, we computed a score of model goodness (the negative free energy as an approximation to the log evidence (Penny, 2012)) and averaged over the posterior means. **Figure 5** illustrates that the conclusions drawn from this analysis do indeed depend on the integration scheme used to perform inference.

**Figure 5.**
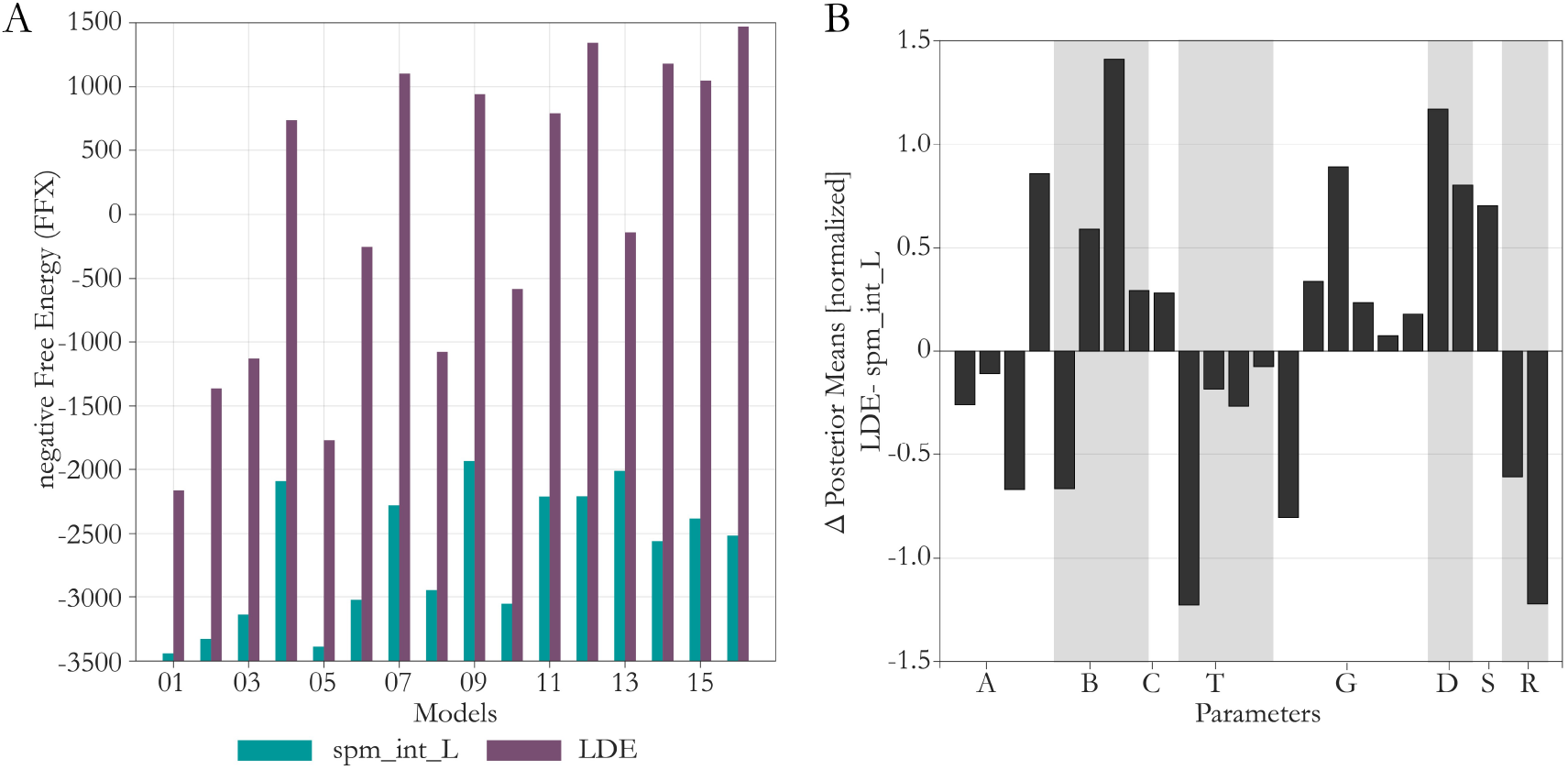
Comparison of model inversion results under different DDE schemes, using empirical data. A) Fixed effects Bayesian model comparison (N=8, higher bars represent higher evidence in favour of a model). B) Difference in posterior means across the two integration schemes (*LDE - spm_int_L*) for model 16. Bars depict normalized difference (t-statistic assuming repeated measures). Parameter labels: Extrinsic Connectivity (A), Modulation (B), Driving Input (C), Kernel Decay (T), Kernel Gain (G), Extrinsic Delays (D), Sigmoid Activation (S), Input Shape (R).

In terms of model selection, we could observe a difference in the selected winning model (**Figure 5A**). The *LDE* integration scheme deems model 16 (i.e. a model including all modulations) the most likely model to have generated the data. By contrast, *spm_int_L* considers model 9 the winning model. In this model, the self-inhibition of the superficial pyramidal cell in region 1 is not modulated by *deviant*. In principle, one can also formally draw a comparison between the two integration schemes, where the *LDE* scheme clearly outperforms *spm_int_L*, in the sense that it achieves a higher negative free energy for each of the models considered. While BMS might not be the most common approach to assess the goodness of an integration scheme, it is useful to provide a holistic perspective as long as all other components (data, generative model, model inversion methods) are identical and only the DDE integration scheme is changed (see also Lemarechal et al., 2018). In particular, given that a difference in the negative free energy >3 is typically considered as decisive evidence for one model versus another, the far greater differences shown in **Figure 5** illustrate that DDE-dependent differences can be of a magnitude which are relevant for the conclusions drawn from model comparison.

In terms of posterior estimates, we observed differences in a number of parameter estimates, in particular modulatory parameters, kernel decays, kernel gain, and delays (**Figure 5B**). The values in (**Figure 5B**) are z-transformed, assuming repeated measures, i.e.

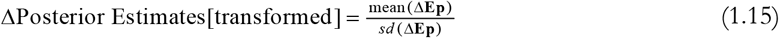

for Δ**Ep** = **Ep**_LDE_ − **Ep**_spm_int_L_, where **Ep** denotes the posterior means^4^.

Additionally, we found a number of sign flips in the estimates of modulatory and delay parameters when comparing the two integration schemes (**Figure 6**). This is a concern when using parameter estimates as readouts of neuromodulatory action since wrongly inferred directionalities of effects (i.e., excitatory vs inhibitory) could lead to significant misinterpretations.

**Figure 6.**
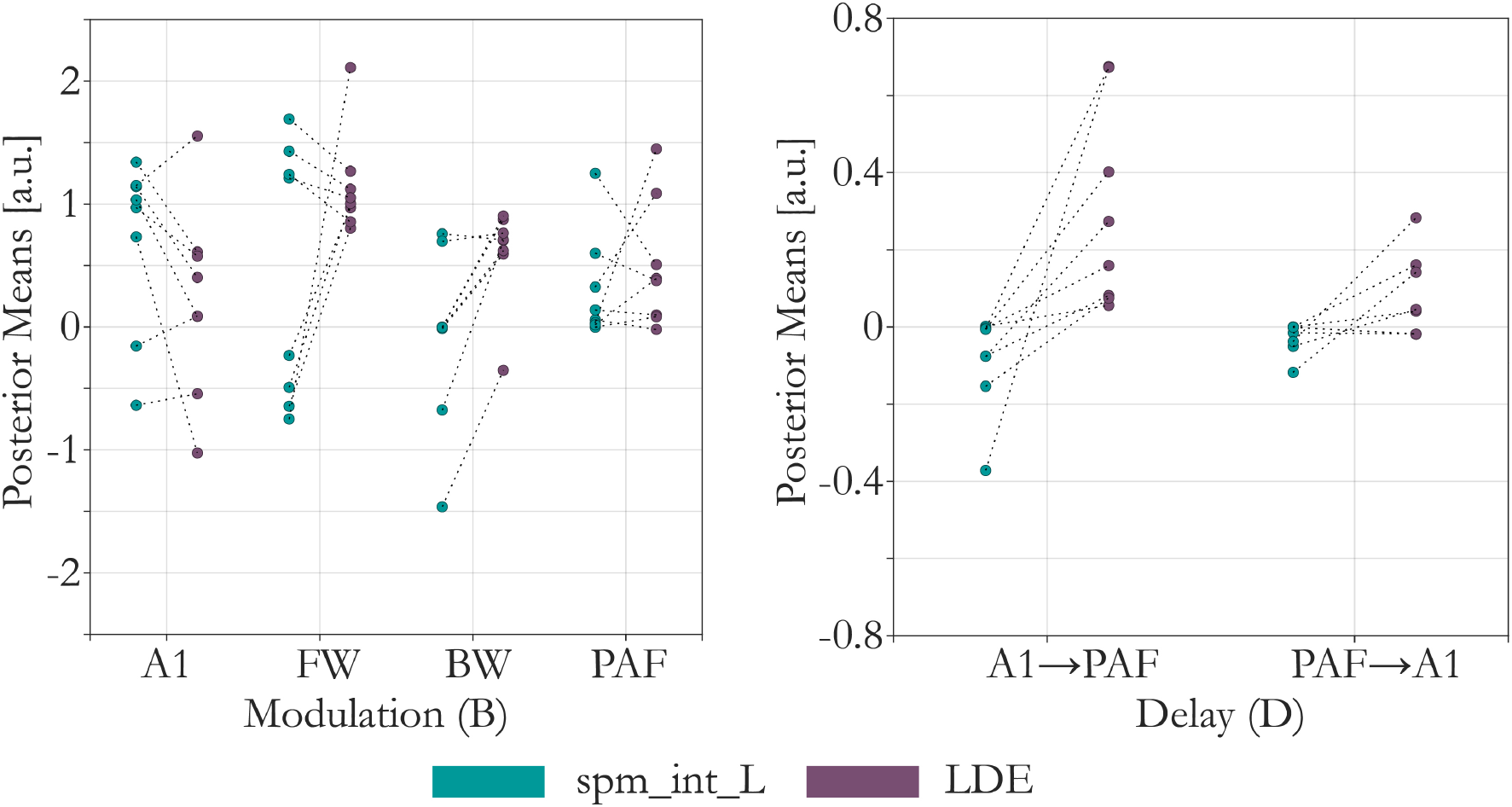
Detailed comparison of parameter estimates regarding modulation and delay. Single dots represent single inversion results. Dotted lines depict the same, underlying data (i.e. animal). Left) Modulation parameter estimates (B, see Figure 5B) for intrinsic modulation in A1 and PAF and modulation of forward (FW) and backward (BW) connections. Right) Delay parameter estimates (D, see Figure 5B) for forward (A1 →PAF) and backward (PAF→A1) delays.

The multistart procedure we used here makes it computationally infeasible to use *dde23* in the same fashion (see the following section on the speed of the integration schemes). In order to evaluate the solutions from *spm_int_L* and *LDE* in comparison to *dde23* in an efficient manner, we used the posterior means computed with *spm_int_L* and *LDE*, and integrated the system with *dde23* using the respective values. **Figure 7** compares the difference in the predictions for two datasets (rodents). Clearly, the prediction for one of the rodents differs vastly between s*pm_int_L* and *dde23 (*Dataset A), in particular with a large difference in the posterior auditory region (PAF). On the other hand, *LDE* seems to produce a very similar prediction to *dde23* for the same parameter values. Both findings are in agreement with the previous simulations (compare **Figure 3**). There is a hint of a small integration artefact visible in *LDE* (Dataset B), which however should be well within the levels of expected (irreducible) noise of the generative model.

**Figure 7.**
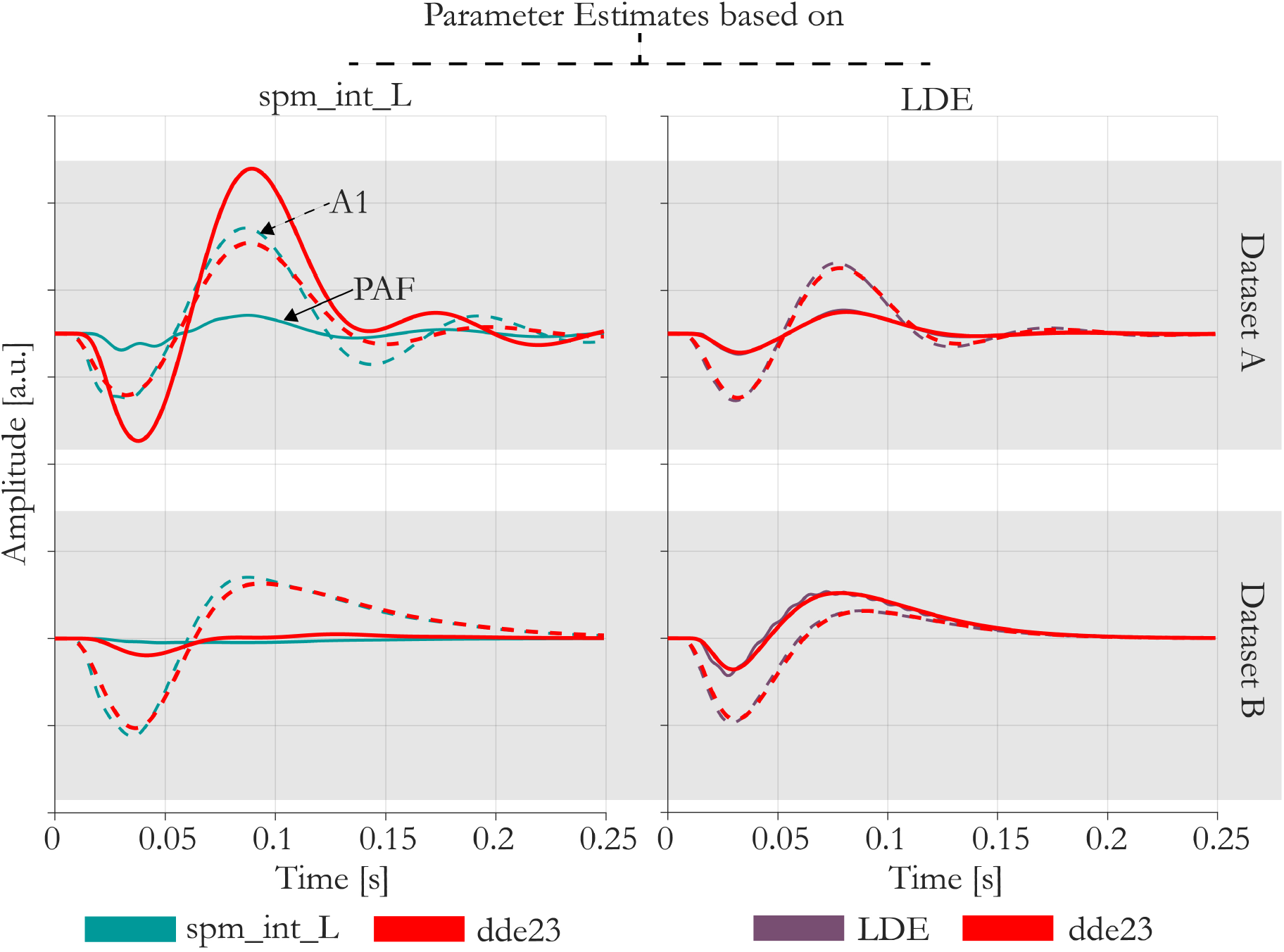
Integrated systems with *dde23* based on the parameter estimates computed using *spm_int_L* and *LDE*. Prediction for two different datasets (rodents) shown for illustration. Dotted lines depict the predicted activity in A1, solid lines depict the predicted activity in PAF (standard condition).

### Speed Comparison

We tested the runtime for all three integration schemes for two different sizes of the networks (**Table 3**). Since none of the integrators was carefully optimized for speed, the numbers below are only meant to provide an approximate comparison. **Table 3** shows that *spm_int_L* is extremely efficient since the only added complexity compared to the non-delayed Ozaki (ODE) integration is the computation of a single matrix inversion. Our continuous extension of the *LDE* method does not substantially increase the runtime. Specifically, for two sources, it is slower than *spm_int_L* by a factor of approximately 2.5, a difference that increases only linearly with the number of regions. By contrast, *dde23* requires much more computational resources. The reason is two-fold. First, *dde23* performs multiple RK steps, i.e. multiple evaluations of the dynamic equation. Secondly, it uses subsampling to control the error, thus requires more integration steps. The latter disadvantage of course is reduced, when one considers subsampling for the other integrators, or time-dependent evaluations of the Jacobian (*spm_int_J*).

**Table 3.** Average integration time for the three types of DDE integrators and a two and four region DCM (18 and 36 states respectively). Signals are integrated over 500 ms (at an integration step size of 1 ms). Averages were computed over 1000 random initialization of the delays.

|  | 2 Sources |  |  | 4 Sources |  |  |
| --- | --- | --- | --- | --- | --- | --- |
|  | *spm_int_L* | *LDE* | *dde23* | *spm_int_L* | *LDE* | *dde23* |
| average integration time (in ms) | 42 | 112 | 1251 | 47 | 239 | 4714 |
| average integration time (in % of *dde23*) | 3.4 | 9 | 100 | 1 | 5.1 | 100 |

DCM analyses of empirical electrophysiological datasets that consider multiple sources (regions) often involve on the order of 40 free parameters (e.g. see (Garrido et al., 2007)). In this case, a single optimization step requires around 41 integrations (one for the prediction, 40 for the gradients). On the other hand, the standard variational Bayesian optimization algorithm in SPM (variational Laplace) requires roughly 50-80 steps to invert a model. If one assumed an additional integration cost of 4 s for *dde23* compared to the other integrators, this would amount to a difference of well over two hours per model, per subject, per condition. In fact, at a relative error tolerance of 0.1%, *dde23* took over six hours for the inversion of a single model consisting of five regions (Lemarechal et al., 2018). In comparison, *spm_int_L* would only require a couple of minutes to perform the full inversion.

## Discussion

An attractive strategy in Translational Neuromodeling and Computational Psychiatry (TN/CP) is the development and clinical use of “computational assays”, i.e. generative models for inferring disease mechanisms from non-invasively obtained neurophysiological and behavioural data (Stephan & Mathys 2014). Biophysically interpretable network models in particular have a promising potential for enabling patient-specific inferences about synaptic dysfunction ((Moran et al., 2011; Murray et al., 2012; Gilbert et al., 2016; Symmonds et al., 2018; Adams et al., 2020); for review, see (Frässle et al., 2018)). A specific class of generative models potentially suitable for this approach are DCMs of electrophysiological data ((David et al., 2006); for review, see (Kiebel et al., 2009)).

One critical condition for turning generative models into tools for routine clinical use is that parameter estimates obtained by these models can be interpreted reliably. This, in turn, requires sufficient robustness of the numerical procedures involved, such as approximate Bayesian inference techniques and the integration of (delay) differential equations. Here, we focused on the latter issue.

In this technical note, we followed up on the previous work by Lemarechal et al. (2018) who demonstrated that the standard integration method used in SPM12 for the integration of the DDEs underlying electrophysiological DCMs might not always have the desired degree of accuracy, with potentially detrimental consequences for parameter estimation and their interpretation in terms of synaptic function. While Lemarechal et al. (2018) provided an alternative integration scheme with high accuracy, based on MATLAB’s *dde23* integrator, this solution is up to two orders of magnitude slower than the default scheme, depending on the desired error tolerance (Lemarechal et al., 2018). It is therefore still an open question how the accuracy of integrating DDEs in DCMs could be improved while avoiding excessively long compute times. In this study, we provide an answer to this question and suggest a novel DDE integrator for DCM – Linearized Delayed Euler (*LDE*) – that is highly accurate and yet computationally efficient.

In a first step, we confirmed the earlier finding by Lemarechal et al. (2018) that the default integration scheme does not always accurately account for delay effects. We demonstrated this by performing simulations using three different dynamical systems with known properties. In these simulations, *spm_int_L* failed to capture delays appropriately, resulting in considerable integration errors if delays became too large in comparison to the intrinsic time constants of the system. For example, *spm_int_L* did not result in the analytically predicted oscillation of a one-dimensional, delayed exponential decay and violated the temporal succession of activity propagation imposed by the delays in a network of coupled harmonic oscillators. By contrast, these problems were not visible when applying the newly developed *LDE* nor *dde23*.

Our results from applying DCM to empirical data using both *spm_int_L* and *LDE* illustrated that conclusions can differ when different DDE schemes are used. Both network identification (assessed through fixed-effects BMS) and parameter estimates differed considerably between *spm_int_L* and *LDE*. Model comparison, in terms of the negative free energy as an approximation to log evidence, indicated that the use of *LDE* greatly improved the performance of the generative model: The difference in negative free energy Δ*F* across the two integration schemes was substantially above standard thresholds used for deciding between competing models (Penny, 2012). In order to compare the solutions from *spm_int_L* and *LDE* against an independent (accurate but slow) integration scheme, we integrated the system with MATLAB’s *dde23* using the posterior means estimated under *spm_int_L* and *LDE*, respectively **Figure** 7). This comparison indicates that *LDE* produces very similar predictions to *dde23* for the same parameter values, whereas marked differences occur under *spm_int_L*. This result is in agreement with the simulation analysis (**Figure 3**).

Given the observed weaknesses of *spm_int_L*, can it be used in practice and when is it safe to do so? Our simulations showed that the answer to this question depends on the relation between delays and the time constants of the system of interest. As shown in **Figure 4**, relevant differences occur for delays as small as approx. 10 % of the time constants of the system. This has implications for the practical application of *spm_int_L* to human EEG/MEG data, the most common use case of electrophysiological DCMs. For example, auditory ERPs typically show frequencies in the range of 10 – 20 Hz, i.e. time constants of 50-100 ms (Haenschel et al., 2000). Conduction delays in human cortex are not known for all types of connections; however, information exists on callosal connections in human cortex whose mean estimates range between approximately 5-10 ms, albeit with considerable variability (Caminiti et al., 2013). Across species, animal studies have reported delays between 0.5 ms and 42 ms for cortico-cortical connections (for a review of existing studies, see (Swadlow and Waxman, 2012)). Thus, expected delays are in a range where differences in the accuracy of integration schemes are likely to be numerically relevant^5^. We emphasise that this does not mean that the use of *spm_int_L* in electrophysiological DCMs will necessarily lead to flawed inferences. Instead, the impact of potential inaccuracies will depend on the specific circuit considered and on the experimental paradigm that perturbs its activity; these factors determine delays and time constants which, in turn, determine which degree of accuracy is needed for the DDE integrator.

As described above (Eqs. 1.3-1.6), *spm_int_L* rests on absorbing a delay operator into the Jacobian. It should be noted that this scheme can be generalised, using a high-order Taylor expansion (Friston et al., 2014). This is particularly important when dealing with fast electrophysiological responses; for example, gamma band activity in induced responses. In this situation, one can use an extension of *spm_int_L*, as described in the appendix of Friston et al. (2014). This high-order approximation ameliorates some of the problems discussed above; however, it comes at a computational cost. In brief, this is because the first-order approximation enables the delay operator to be computed analytically. However, when one includes higher-order terms, one has to find the roots of a matrix polynomial (e.g., using a Robbins-Munro algorithm). Typically, when integrating electrophysiological systems that show fast fluctuations, one needs to increase the order of the delay approximation considerably (the default is N=256 in *spm_dcm_delay* in SPM) (Karl Friston, personal communication). For smoothly varying evoked responses in the alpha range, the first-order (N=1) approximation can be sufficient – although, our analyses suggest that an *LDE* would be preferred because it is more robust.

In order to avoid the necessity for a case-by-case analysis and resolve uncertainty, it is desirable that a DDE integration method is in place that can be used universally. The sophisticated *dde23* DDE integrator in MATLAB represents one option; however, its poor computational efficiency renders it a suboptimal choice for many practical applications of DCM.

Here, we proposed and implemented an alternative DDE integration scheme based on the principle of continuous extension of ODE methods (Feldstein, 1964). Across all simulations, the new *LDE* integrator respected the nature of delayed effects at reasonable step size. Importantly, this did not lead to a marked increase in computation time (**Table 3**). While this paper focuses on the application of *LDE* to the convolution-based formalism of DCMs, the new integration scheme can be equally applied to more advanced formulations of DCMs (e.g. conductance based models). It is also notable that our scheme could in principle accommodate different update steps (e.g. Euler, Ozaki, Runge-Kutta, etc.) and can thus be extended flexibly. Having said this, even with the simple Euler formulation described in this article, it performed well in comparison to the reference scheme (*dde23*). To facilitate the use of this method, the MATLAB code has been made available that integrates *LDE* smoothly into the existing functionality of SPM12 and is publicly available as part of the open-source software collection TAPAS (https://www.translationalneuromodeling.org/tapas).

We hope that the availability of this method and its compatibility with the SPM functions for DCMs will facilitate a wide application in the community, help avoid integrator-dependent confounds in the application of DCMs for EEG/MEG, and generally contribute to the development of computational assays in TN/CP.

## Authors’ contributions

**Dario Schöbi:** Conceptualization, Methodology, Software, Formal Analysis, Writing – Original Draft **Cao Tri Do:** Software, Validation, Writing – Review & Editing **Stefan Frässle:** Validation, Writing – Review & Editing **Marc Tittgemeyer:** Resources, Writing – Review & Editing **Jakob Heinzle:** Conceptualization, Methodology, Supervision, Writing – Review & Editing **Klaas Enno Stephan:** Resources, Conceptualization, Supervision, Writing – Review & Editing; Funding Acquisition

## Acknowledgements

We would like to thank Karl Friston for helpful comments on the manuscript. This work was supported by the René and Susanne Braginsky Foundation (KES) and the University of Zurich (KES).

## Footnotes

1 The methods and results presented here are based on the PhD thesis by Dario Schöbi at ETH Zürich (Schöbi D (2020) Dynamic causal models for inference on neuromodulatory processes in neural circuits. In: ETH Zurich.).

2 Here, c*ritical* must be understood in a qualitative fashion for when the linear approximation becomes inaccurate. Critical magnitudes for delays are most likely a function of the integration step size and the frequencies of the system.

3 To avoid the risk of comparing suboptimal solutions due to local extrema, computational optimization of the models was performed starting from 100 starting values (i.e. a multistart procedure). All results presented correspond to the solution based on the starting value resulting in the highest negative free energy (also see Schöbi D (2020) Dynamic causal models for inference on neuromodulatory processes in neural circuits. In: ETH Zurich.)

4 For *N* = 8, normalized values of around z = 0.83 correspond to p<0.05 (uncorrected).

5 The current prior in SPM12 expects conductance delays between cortical columns on the order of 9-25 ms (i.e. approximately 95% of the prior probability mass).

## Notes

### Competing Interest Statement

The authors have declared no competing interest.

